# How to design optimal eDNA sampling strategies for biomonitoring in river networks

**DOI:** 10.1101/2020.05.18.102368

**Authors:** Luca Carraro, Julian B. Stauffer, Florian Altermatt

## Abstract

The current biodiversity crisis calls for appropriate and timely methods to assess state and change of biodiversity. In this respect, environmental DNA (eDNA) is a highly promising tool, especially for aquatic ecosystems. While initial eDNA studies assessed biodiversity at a few sites, technology now allows analyses of samples from many points at a time. However, the selection of these sites has been mostly motivated on an ad-hoc basis, and it is unclear where to position sampling sites in a river network to most effectively sample biodiversity. To this end, hydrology-based models might offer a unique guidance on where to sample eDNA to reconstruct the spatial patterns of taxon density based on eDNA data collected across a watershed.

Here, we performed computer simulations to identify best-practice criteria for the choice of positioning of eDNA sampling sites in river networks. To do so, we combined a hydrology-based eDNA transport model with a virtual river network reproducing the scaling features of real rivers. In particular, we conducted simulations investigating scenarios of different number and location of eDNA sampling sites in a riverine network, different spatial taxon distributions, and different eDNA measurement errors.

We identified best practices for sampling site selection for taxa that have a scattered versus an even distribution across the network. We observed that, due to hydrological controls, non-uniform patterns of eDNA concentration arise even if the taxon distribution is uniform and decay is neglected. We also found that uncertainties in eDNA concentration estimates do not necessarily hamper model predictions. Knowledge of eDNA decay rates improves model predictions, highlighting the need for empirical estimates of these rates under relevant environmental conditions. Our simulations help define strategies for the design of eDNA sampling campaigns in river networks, and can guide the sampling effort of field ecologists and environmental authorities.

## 1 Introduction

The recently released report by the Intergovernmental Science-Policy Platform on Biodiversity and Ecosystem Services (IPBES) shows that global biodiversity is declining in an unprecedented way, and effective societal and policy responses are needed now more than ever [Díaz et al., 2020]. While all ecosystems are affected, showing both strong declines in biodiversity and associated ecosystem functions, freshwater and riverine ecosystems in particular are among the most concerned. Changes in land use, climate, damming and hydropower, and chemical pollution heavily impair riverine ecosystems [Darwall et al., 2018; Reid et al., 2019; Vörösmarty et al., 2010], such that their status is a matter of primary societal and political concern [Dudgeon, 2019]. These changes call for a rapid understanding and documentation of the state but also change of biodiversity. Adequate strategies for freshwater biodiversity preservation must thus adopt efficient monitoring tools, and comprise strategies that acknowledge the characteristic spatial structure of river networks and their biodiversity [Altermatt, 2013]. In this context, the use of environmental DNA (eDNA) constituted a major breakthrough of the last decade [Beng and Corlett, 2020; Bohmann et al., 2014; Ficetola et al., 2008; Kelly et al., 2014; Thomsen and Willerslev, 2015]. The eDNA technique consists in the detection of organisms based on their DNA extracted from environmental samples (e.g. from soil or water) [Taberlet et al., 2012], and can cover organisms ranging from bacteria to eukaryotes. Compared to traditional species sampling methods, eDNA has the advantage of being minimally invasive and fast, yet able to detect multiple species thanks to metabarcoding [Deiner et al., 2017; Pawlowski et al., 2018], including rare and elusive species [Jerde et al., 2011; Mächler et al., 2014]. This gives the potential for biodiversity assessments at high spatial and temporal resolutions [Altermatt et al., 2020; Bohmann et al., 2014; Pawlowski et al., 2018].

A key feature of such a spatially and temporally highly resolved biodiversity assessment is maximizing the quantity and quality of the data gathered with as little effort as possible. While initial studies on biodiversity assessments in riverine ecosystems sampled at one or few locations within a single watershed, without the goal of a catchment-level perspective (e.g. Mächler et al. [2014]; Thomsen et al. [2012]), more recent works with a biodiversity focus aimed at resolving diversity across the whole catchment [Deiner et al., 2016; Mächler et al., 2019; Sales et al., 2020]. This immediately brings up the question of where to sample, and how many samples to take, to effectively assess biodiversity across a catchment. The costs of sampling and processing of eDNA may scale much less than linearly, both in terms of financial and time costs [Sengupta et al., 2019]; however, to maximize the efficiency of eDNA, optimizing sampling strategies is paramount. Guidelines on where to best take eDNA samples in river networks are thus needed, and these guidelines must consider the origin, transport and decay of eDNA along the waterway.

Research on molecular and bioinformatic aspects of eDNA analyses has massively expanded within the last few years [Alberdi et al., 2017; Beng and Corlett, 2020; Calderón-Sanou et al., 2019; Deiner et al., 2017; Ficetola et al., 2016; Garlapati et al., 2019; Leray et al., 2019]. Much less focus, however, has been attributed to the “ecology of eDNA”, namely “its origin, state, transport, and fate within the environment” [Barnes and Turner, 2015], and how these factors influence its detection [Barnes and Turner, 2015; Harrison et al., 2019]. Indeed, in riverine environments, hydrological transport of material containing genetic information makes eDNA a carrier of information on biodiversity of the upstream catchment [Deiner et al., 2016]. While this fact underlines the crucial role of eDNA as a tool to monitor biodiversity at large scales, it also gives rise to further challenges with respect to the reconstruction of spatial patterns of biodiversity. Essentially, the eDNA sampled at a river’s cross-section results from the aggregation of the dynamics of particle transport from a number of upstream sources (i.e. the locations of the target species) along a dendritic river network towards the sampling site. Importantly, eDNA advection is subject to decay processes typically dependent on several abiotic and biotic factors, as well as on hydrological conditions [Barnes et al., 2014; Shogren et al., 2017; Strickler et al., 2015], resulting in downstream travelling distances ranging from meters to tens or hundreds of kilometers [Deiner and Altermatt, 2014; Jane et al., 2015; Pont et al., 2018; Shogren et al., 2017]. Thus, it is clear that comprehensive eDNA studies in riverine environments cannot ignore hydrological and geomorphological concepts. Research at the boundary between hydrology and molecular ecology is needed, as the sole focus on the molecular aspects of eDNA analyses would lead to an incomplete application of the method and undermine its power as a biomonitoring tool.

Recently, physically-based models with different degrees of complexity have been developed in order to assess dynamics of transport and decay of eDNA in water. Sassoubre et al. [2016] applied a simple mass-balance model to a tank experiment; Shogren et al. [2016] studied eDNA transport in a column experiment; Shogren et al. [2017] used a simple transport model in a flume experiment; Nukazawa et al. [2018] and Sansom and Sassoubre [2017] formulated 1D advection models in river stretches; Andruszkiewicz et al. [2019] and Fukaya et al. [2018] applied 3D advection-diffusion equations to study eDNA transport in marine bays. Carraro et al. [2017, 2018] coupled a first-order formulation of eDNA decay with well-established knowledge on the geomorphology and hydraulic properties of river networks [Leopold and Maddock, 1953; Rodriguez-Iturbe and Rinaldo, 2001] to infer the upstream distribution of target species based on eDNA data sampled at multiple locations across the river network. The ability of such types of models to accurately predict spatial patterns of biodiversity substantially relies on the accuracy of the eDNA measurements. However, literature on an optimal layout of eDNA sampling strategies is rather poor: Dickie et al. [2018] reviewed practices of sampling protocols in terrestrial and freshwater eDNA studies, but did not specifically address the issue of positioning eDNA sampling sites across a river network; Bylemans et al. [2018] studied the effect of sampling intensity and replication at fixed sampling sites on the estimation of fish biodiversity in a river, and found that a larger number of replicates at downstream sites is needed to fully assess biodiversity; Wood et al. [2020] observed that, due to transverse hydrodynamic dispersion, detection rates are maximized if eDNA is collected at some distance downstream of a known source, otherwise the dispersion plume of eDNA would likely be missed.

No studies to date, however, have investigated the optimal positioning of sampling sites across whole catchments, although such information would be highly relevant to the effectiveness of eDNA sampling campaigns in rivers. Here, we fill this gap by making use of computer simulations in order to assess different scenarios of eDNA release, transport and detection. We test these scenarios based on the application of the hydrology-based eDNA transport model of Carraro et al. [2018] using synthetic analogues of river networks (so-called Optimal Channel Networks, see Carraro et al. [2020]; Rinaldo et al. [2014]). This allows testing realistic spatial scenarios in a generic setting that is not constrained to the particular shape of a real river network. In the studied scenarios, we explored varying numbers and locations of the sampling sites, spatial distributions of the target species as a proxy of eDNA release, and assumptions on decay rate and eDNA measurement errors. Together, this allowed the identification of generalizable rules of optimal eDNA sampling strategies in rivers.

## 2 Materials and methods

This section is structured as follows: first, we introduce the tools needed for the subsequent computer simulations, namely an eDNA transport model and a virtual river network; second, we show how such tools are used to assess patterns of eDNA concentration across a river produced by particular spatial distributions of taxon density and decay rate values; third, we describe the strategy and details of the computer simulations aimed at assessing optimal eDNA sampling strategies. Note that, as this section substantially relies on mathematical formulations, we introduce every subsection with a brief description in which the relevant information is outlined, then followed by the more technical details. All mathematical symbols used in the following are listed in Table S1.

### 2.1 The eDITH model

To simulate transport of eDNA in the water across a river network, we utilize the approach of Carraro et al. [2018], subsequently referred to as eDITH (*eD*NA *I*ntegrating *T*ransport and *H*ydrology). This approach exploits knowledge on hydrology and geomorphology of river networks to transform a map of eDNA production *p* across a catchment into a map of eDNA concentration *C* in stream water. As a first approximation, *p* can fairly be assumed to be proportional to relative taxon density (see Carraro et al. [2018] for a discussion on caveats of this assumption), which enables considering the eDITH model as a function that relates a spatial distribution of taxon density to a spatial distribution of eDNA concentration. Note that we do not make specific assumptions on the molecular state of the DNA extracted from the environments; we only assume that organisms can be detected based on their DNA in environmental samples, and that this DNA signal is subject to generic processes describing production, transport and decay.

Notably, the eDITH model can be used in two different ways. First, assuming that the spatial distribution of a taxon is known, it allows transforming such distribution into the corresponding pattern of eDNA concentration. Second, eDITH can be used in an inverse modelling approach, where eDNA concentrations at some locations within the river network are measured, and the model is used to infer what is the pattern of *p* that is most likely to have generated such values of eDNA concentration. In the following application, both uses of the eDITH model will be employed.

Note that all parameters involved in the relationship between *C* and *p* (except the decay rate) relate to the morphology of the river network, and can thus be assessed with sufficient precision with a combination of GIS-based river network extraction, field observations and power-law scaling relationships for the hydraulic variables of interest [Leopold and Maddock, 1953; O’Callaghan and Mark, 1984; Rodriguez-Iturbe and Rinaldo, 2001]. The value of the eDNA decay rate can be directly inferred from model calibration; alternatively, specific information on eDNA decay dynamics (if available) could be used to define such value (or, at a minimum, a range thereof), thus likely improving the model’s prediction skill.

Technical details of the eDITH model are hereafter provided. Let us consider a river network discretized into *N* nodes. For a given taxon, the eDNA concentration *C_j_* at node *j* can be expressed as:

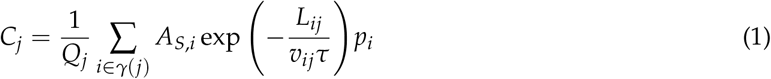

where *Q_j_* is a characteristic value of water discharge at node *j*; *γ*(*j*) identifies the set of nodes upstream of *j*; *A_S_*,*_i_* is the source area (i.e. the habitat extent) of node *i*; *p_i_* is the eDNA production rate at node *i*; exp [−*L_ij_*/(*v_ij_τ*)] is a first-order exponential decay factor, in which *L_ij_* is the length of the path between *i* and *j*, *v_ij_* the average water velocity along this path, and *τ* a characteristic decay time for the taxon’s eDNA in running water (i.e. inverse of a decay rate, which accounts for all sources of eDNA depletion, from degradation to gravity-induced deposition). For a water-dwelling taxon (the case that we investigate in this application), *A_S_*,*_i_* can be seen as the water surface area at node *i*: *A_S_*,*_i_* = *L_i_w_i_*, where *L_i_* and *w_i_* are the length and width of the river stretch corresponding to node *i*, respectively.

### 2.2 Optimal Channel Networks

The virtual river network used in the following simulations is an Optimal Channel Network (OCN) built via the R-package *OCNet* [Carraro et al., 2020]. OCNs are idealized constructs that reproduce the topological and scaling features of real river networks, and are therefore suitable for simulation studies on various ecological and ecohydrological issues [Rinaldo et al., 2014; Rodriguez-Iturbe et al., 1992]. In this application, we analyzed the outputs of the eDITH model on an OCN that represents a catchment covering an area of 400 km^2^ (Figure 1).

**Figure 1:**
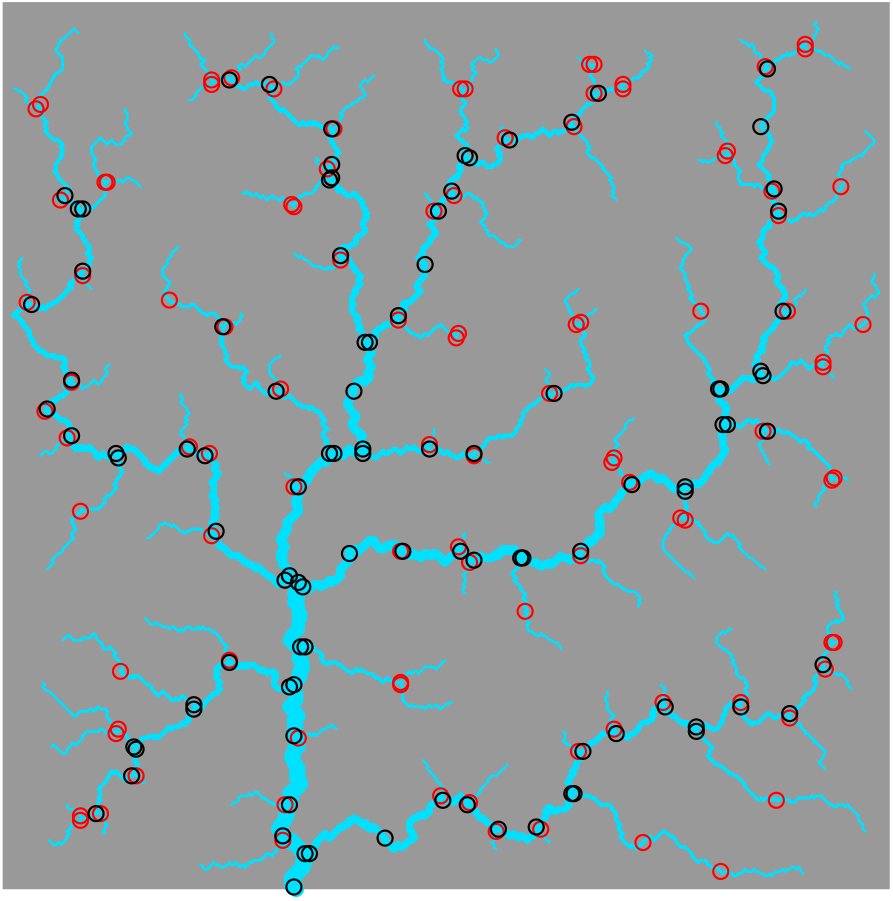
Representation of the optimal channel network used in this study, spanning a square of side 20 km. The aggregation of the OCN at the AG level (see Carraro et al. [2020]) identifies 200 nodes, which are here displayed at the downstream end of the corresponding river reaches. Red indicates nodes whose drainage area is lower than the median drainage area across the 200 nodes (referred to in the text as upstream nodes); black identifies nodes with drainage area higher than the median (downstream nodes).

The OCN was built on a square lattice whose side is made up of 400 pixels, each of which is assumed to represent a 50-m by 50-m cell. A threshold area of 461 pixels (*A_t_* = 461 · 0.05^2^ = 1.15 km^2^) was used to distinguish the fraction of lattice pixels that effectively constitute the river network. At this so-called RN (river network) aggregation level, the network is constituted by 4468 nodes (each corresponding to a 50-m pixel with drainage area ≥ *A_T_*; see Carraro et al. [2020] for details on aggregation levels of an OCN). Subsequently, such network was further coarsened into the AG (aggregated) level, where nodes represent sources and confluences of the network at the RN level, while edges follow the drainage directions previously identified. Additional nodes were added in order to split the edges into portions not longer than 2.5 km (option *maxReachLength* in function *aggregate_OCN* of *OCNet*). Such maximum length value was arbitrarily imposed in order to partition the OCN into reaches of limited size, where abiotic (water discharge and velocity) and biotic (taxon density) variables could fairly be considered homogeneous. The resulting number of nodes at the AG level is 201. Note that, since all nodes except the outlet node are associated to an edge directed downstream, the river network at the AG level is essentially partitioned into 200 segments, while the outlet node is immaterial. The choice of *A_T_* was purposely operated to obtain such result. According to a well-established approach in hydrology [Leopold and Maddock, 1953; Rodriguez-Iturbe and Rinaldo, 2001], water discharge *Q*, river width *w* and water velocity *v* across the whole river network were calculated via power-law functions of drainage area *A*:

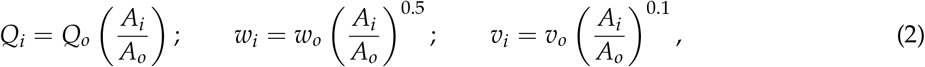

where subscript *o* identifies the outlet node. The values specified at the outlet node were *Q_o_* = 10 m^3^s^−1^, *w_o_* = 10 m, *v_o_* = 1 ms^−1^, *A_o_* = 400 km^2^.

### 2.3 Analysis of eDNA concentration patterns across a river network

A first set of simulations aimed at qualitatively assessing patterns of eDNA concentrations across the OCN produced by some peculiar distributions of taxon density (expressed as eDNA production rates *p*) and different values of the decay time *τ* (fast decay: *τ* = 1 h; intermediate decay: *τ* = 4 h; no decay: *τ* → ∞). We wanted to compare the eDNA concentration patterns resulting from the two most distinct distributions of taxon density: on the one hand, a uniform distribution, in which the density (i.e. biomass per unit habitat area) of a taxon is evenly distributed across the whole river network, thereby representative of a generalist species; on the other hand, a distribution related to a unique point source, in which the taxon is only present at the node that is farthest from the outlet. The latter represents a rare species, specialized to headwater streams. Indeed, any other spatial pattern of taxon density can be seen as a combination of these two distributions. The uniform distribution is obtained by setting *p_i_* = 1 mol/(m^2^· s) ∀ *i* = 1, …, *N*. The unit for *p* expresses the fact that production rates are amounts of substance (i.e. DNA) per unit habitat area and unit of time [Carraro et al., 2018]. However, units are immaterial in our application, thus we will hereafter refer to a normalized production rate 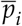, which is equal to 1 ∀ *i* = 1, …, *N* in the uniform distribution case. To enable comparison with the uniform case, the point source distribution is defined by:

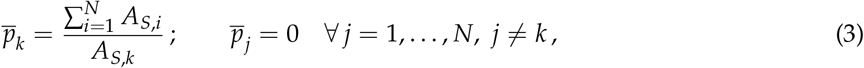

where *k* identifies the node that is farthest from the outlet. Eq. (3) assumes that all the amount of eDNA (per time unit) produced across the river network in the uniform case is now located in node *k*.

In an analogous way, the eDNA concentrations resulting from the application of the eDITH model to these spatial distributions of 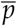 are also normalized in order to facilitate comparison. Normalized concentrations 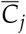 are calculated as:

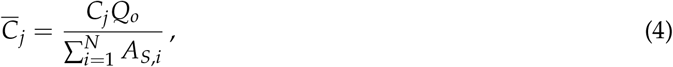

where 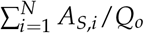 is the concentration that would be measured at the outlet under the hypothesis of no eDNA decay (*τ* → ∞ in Eq. (1)).

In this first exercise, we make use of the refined OCN aggregated at the RN level (*N* = 4468).

### 2.4 Assessing the effect of sampling strategy

#### 2.4.1 Overview

In a second phase, we combined the tools hitherto presented in order to assess the effectiveness of different eDNA sampling strategies across a river network. An overview of the sequence of performed operations is shown in Figure 2. The first step consists in building maps of (normalized) taxon density 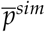 that potentially resemble realistic density maps. Second, we run the eDITH model (Eqs. (1) and (4)) on these maps and yield patterns of (normalized) eDNA concentration 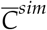. Third, we formulate different sampling strategies in terms of intensity and positioning of sampling sites across the river basin; we then randomly sample sites (i.e. nodes of the network) according to a given strategy, and assume to “observe” the concentration 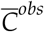 therein (i.e. equal to 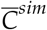 as calculated in step 2, possibly with a measurement error). Fourth, we fit the eDITH model on these observed concentrations, and find a predicted pattern of eDNA production 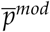 that best reproduces the observed concentrations. Fifth, we compare the resulting maps 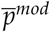 with the original solution 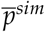 generated in step 1, and assess the prediction skill of the model. All steps are repeated a sufficient number of times to account for the various sources of stochasticity. We adopted a full factorial design, whose factors are listed in Table 1. The total number of simulations performed is 4,500. Details on the factors and levels are hereafter provided. In order to reduce the dimensionality of the problem, this analysis is performed on the OCN aggregated at the AG level (*N* = 200).

**Table 1:**
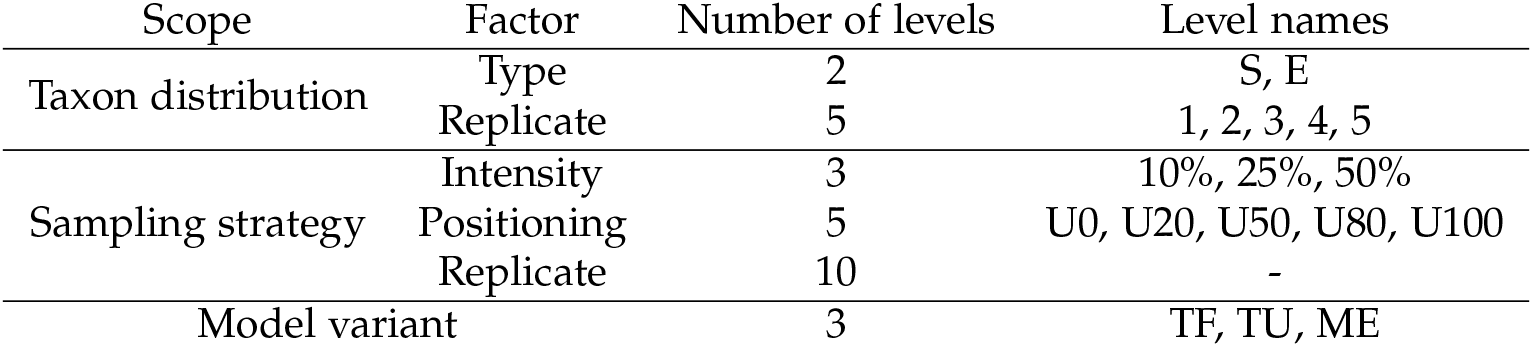
Factors and levels of the full factorial design used in the computer simulations.

**Figure 2:**
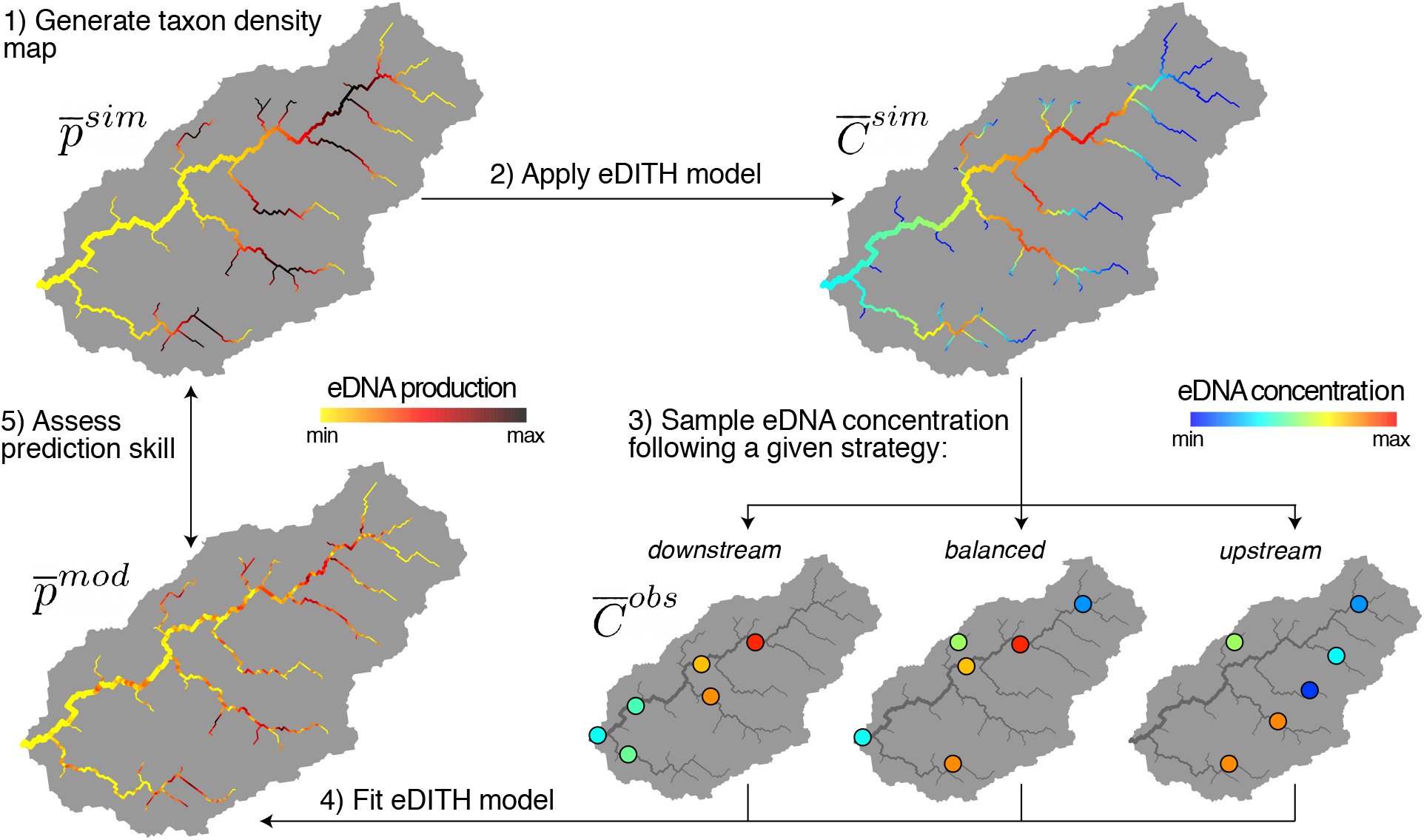
Overview of the computer simulations. In step 1, taxon density maps 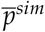 are generated; the example shows a taxon that is abundant only within a narrow elevational range. In step 2, by applying the eDITH model with *τ* = 4 h, corresponding patterns of eDNA concentration 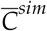 are evaluated. In step 3, following a given sampling strategy (number and location of sampling sites; here, for the same number of sampling sites, three strategies, characterized by differential preference for downstream or upstream reaches, are shown), 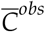 are observed at the sampling locations. In step 4, the eDITH model is fitted on 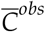, yielding to a modelled taxon distribution 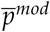. The latter is compared with the original 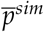 in step 5 to assess the prediction skill of the sampling strategy.

#### 2.4.2 Generation of maps of taxon density

The taxon distribution maps previously introduced (i.e. uniform density versus point source located at the farthest headwater) represent idealized constructs that were introduced for illustrative purposes, although they might be only partially representative of a real taxon distribution. Here instead, we generate more realistic taxon density maps (see e.g. Alther and Altermatt [2018]; Besemer et al. [2013]; Kaelin and Altermatt [2016]; Little et al. [2020]) characterized by a certain number of hotspots across the network, where the taxon is most abundantly located. In particular, we will focus on two types of distributions: a first type, mimicking the distribution of a taxon with scattered distribution (termed S), with few hotspots where the taxon density is high; a second type (E), representing the distribution of a more evenly distributed taxon, with many hotspots in which the taxon density is lower as compared to the previous case.

Taxon density maps are generated by randomly sampling (without replacement) *N_H_* hotspot nodes among the *N* nodes constituting the OCN. On these hotspot nodes, a temporary value of density 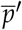 equal to *x* is attributed, while all other network nodes are initially attributed 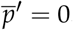 = 0. Next, all nodes immediately downstream or upstream of the hotspot nodes are attributed an additional *x*/2 density (summed to 0 if that node had not been selected, or to *x* or *x*/2 otherwise). The so-obtained temporary densities 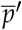 are transformed into a normalized density map via:

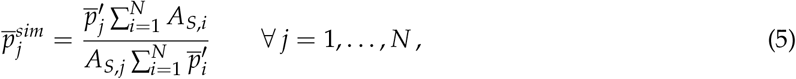

which satisfies the normalization constraint 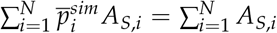 (as in Eq. (3)). For the scattered (S) distribution type, we set *N_H_* = 5, while for type E we set *N_H_* = 50. For each of these distribution types, five different maps are built (see Figure 3).

**Figure 3:**
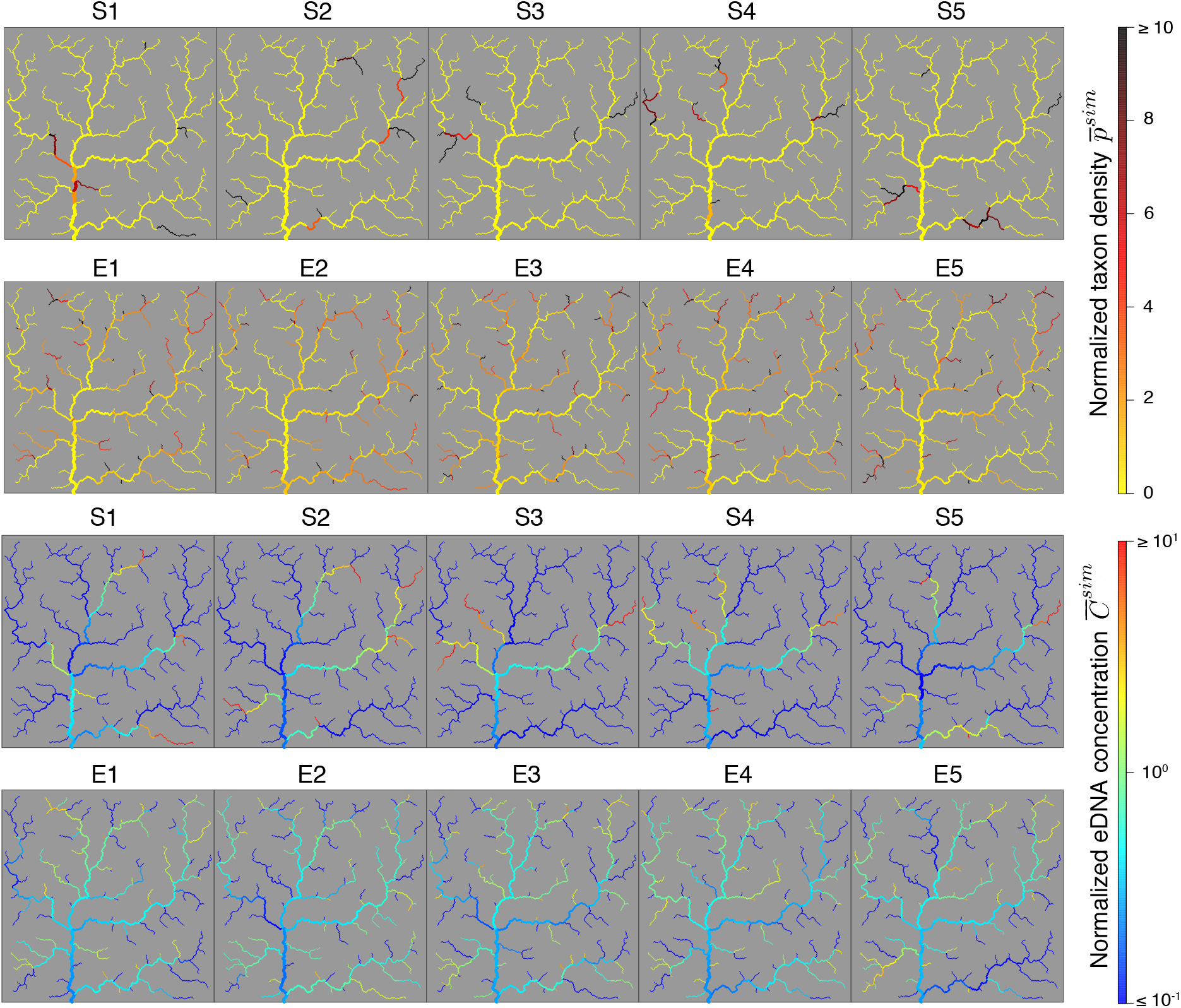
Overview of the ten different maps of normalized taxon density 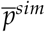 and the corresponding normalized eDNA concentrations 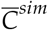 generated (in step 1 and 2 of Figure 2, respectively).

In step 2 of of the computer simulations, we generated patterns of 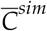 based on the previously derived taxon distribution maps 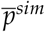 and by fixing *τ* = 4 h. Note that we assumed that sampling sites be located at the downstream end of the river reach to which they are associated. As a result, if *i* is a headwater node (i.e. *γ*(*i*) = {*i*}), the normalized concentration at this site is given by 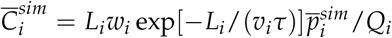 Maps of simulated eDNA concentration patterns are shown in Figure 3.

#### 2.4.3 Sampling strategies

Three different levels of sampling intensity (i.e. number of sampled nodes, where 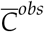 is observed) are adopted: 10%, 25% and 50% of the available network nodes, corresponding to 20, 50 and 100 sampling sites, respectively. For a given level of sampling intensity, five different schemes for sites’ location are investigated. These are defined with the acronyms U0, U20, U50, U80, U100, where the number identifies the percentage of sites that are positioned in the upstream half of the catchment (i.e. among the network nodes whose drainage area *A* is lower than the median of *A*; see Figure 1), while the remaining fraction is sampled among the downstream network nodes. For example, the sampling strategy 10%-U20 consists of 20 sampling nodes, where one fifth of them (4 nodes) are picked from the upstream half of the catchment, and the remaining 16 are chosen within the downstream half. For each sampling strategy, 10 replicated sets of sampling nodes are generated, resulting in a total of 150 sets of sites. Note that the 10 replicates corresponding to strategies 50%-U0 and 50%-U100 actually constitute repetitions of the same set of sampling nodes (the 100 nodes depicted in Figure 1 in black and red, respectively).

#### 2.4.4 Model variants and fitting

We then made different assumptions with respect to the quality of information available on eDNA concentration values and the decay dynamics thereof. In particular, we formulated different model variants, in which we distinguished cases where the eDNA decay rate is known or unknown, and cases where eDNA concentrations at the sampling sites are observed with or without measurement errors (including non-detections).

In a first model variant (termed TF, “tau fixed”), we assumed that, in step 4 (model fitting), *τ* is known to be equal to 4 h, and therefore the only unknown parameters are the 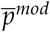 values across the 200 network nodes. Moreover, no measurement error was assumed, hence 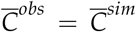 at the sampling sites. In a second model variant (TU, “tau unknown”), we also included *τ* among the unknown parameters, while maintaining 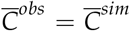. A third model variant (ME, “measurement error”) was obtained by perturbing 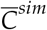 by accounting for a probability of non-detection (as in Carraro et al. [2018]) and a measurement error. In particular, we calculated 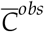 as

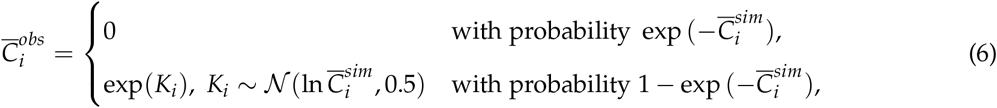

Eq. (6) states that low concentrations 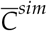 are more likely to result in non-detections: for instance, if 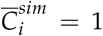, there is a exp(−1) ≈ 36.7% chance to not detect eDNA at site *i*. Moreover, it assumes that observed concentrations 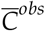 are lognormally distributed around the values 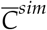 calculated in step 2. The choice of the lognormal distribution is justified by the fact that such distribution does not allow false positives (i.e. if 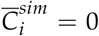 then it must be 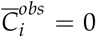). We are aware that false positives in eDNA detection might arise due to e.g. contamination [Ficetola et al., 2016], but we deemed this circumstance beyond the scope of our work. Moreover, in model variant ME, *τ* is assumed to be unknown in the fitting process.

We fitted the models by assuming independent and identically distributed Gaussian errors between modelled 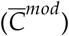 and observed 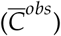 concentrations. Evidence suggests that eDNA concentration in the field might follow a lognormal distribution [Fukaya et al., 2018; Jo et al., 2017]; however, we opted for a normal distribution because it is also directly applicable to null eDNA concentrations, which is not the case for the lognormal distribution. For all three model variants, the so-obtained log-likelihood was maximized by means of the *optimParallel* R-package [Gerber and Furrer, 2019]. All 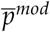 values were constrained between 0 and 250; for models TU and ME, *τ* was constrained between 1 and 251 h. By setting a lower bound of 1 h for *τ*, we forced the calibration algorithm to consider downstream transportation of eDNA when estimating the 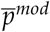 parameters. Failing to do so would often lead to model estimates of *τ* close to 0 and, as a result, totally arbitrary values of 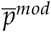 in the unsampled sites, because they would have no impact in the log-likelihood (i.e. all upstream eDNA contribution would be totally degraded before reaching any sampling site).

#### 2.4.5 Assessment of prediction skill

The last step of the computer simulation consists in comparing the estimated map of taxon distribution 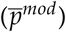 with the original pattern 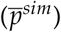 generated in step 1. To do so, we formulated two different criteria: the first one focuses on predictions of presence and absence at the various river reaches, while the second compares values of taxon density.

The first criterion (termed *PA*) expresses the fraction of river reaches where presence or absence were correctly predicted. Since values of 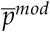 are always larger than zero due to numerical approximations of the model fitting process, we need to impose threshold values in order to convert a map of relative taxon density into a map of occurrences. In particular, we attribute the threshold 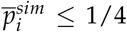 for absence and 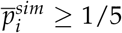 for presence; the small overlap allows e.g. 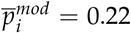 to be a correct estimate of 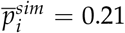. PA reads then:

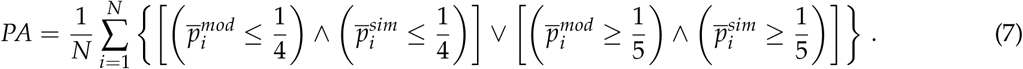

where {*A*} is to be intended as an indicator function, namely equal to 1 if event *A* is true, and null otherwise.

The second criterion (termed *D*) expresses the fraction of river reaches where the modelled taxon density 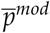 is deemed a good estimate of 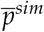:

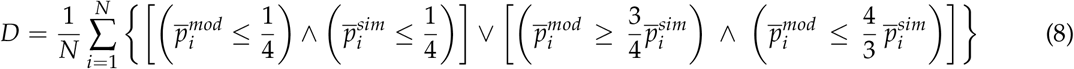

In other words, according to this criterion, 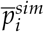 is correctly predicted if the corresponding 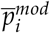 is between 75% and 133% of its value; moreover, if 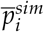 is lower than or equal to 1/4 (hence presumably indicating absence of the taxon in that node), then it must also be 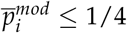. A graphical representation of the two criteria is provided in Figure S1.

## 3 Results

### 3.1 Analysis of eDNA concentration patterns across a river network

eDNA concentration patterns across the OCN as a function of different taxon distributions and decay time values are presented in Figure 4. Remarkably, when the normalized taxon distribution 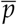 is uniform and no eDNA decay occurs (*τ* → ∞), the resulting eDNA concentration pattern (Figure 4a) is not uniform but rather increases downstream, and reaches its maximum value 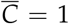 at the outlet (as expected from Eq. (4)). This result can intuitively be explained by the fact that a portion of habitat of unit stream length (in our case, a pixel) located downstream is characterized by a larger width than an analogous portion of habitat located upstream. Hence, uniform taxon density implies higher abundance, and in turn, higher amount of DNA shed at the downstream portion of habitat. As a result, in this scenario eDNA concentration increases in the downstream direction. A more formal explanation is provided in the Supporting Information.

**Figure 4:**
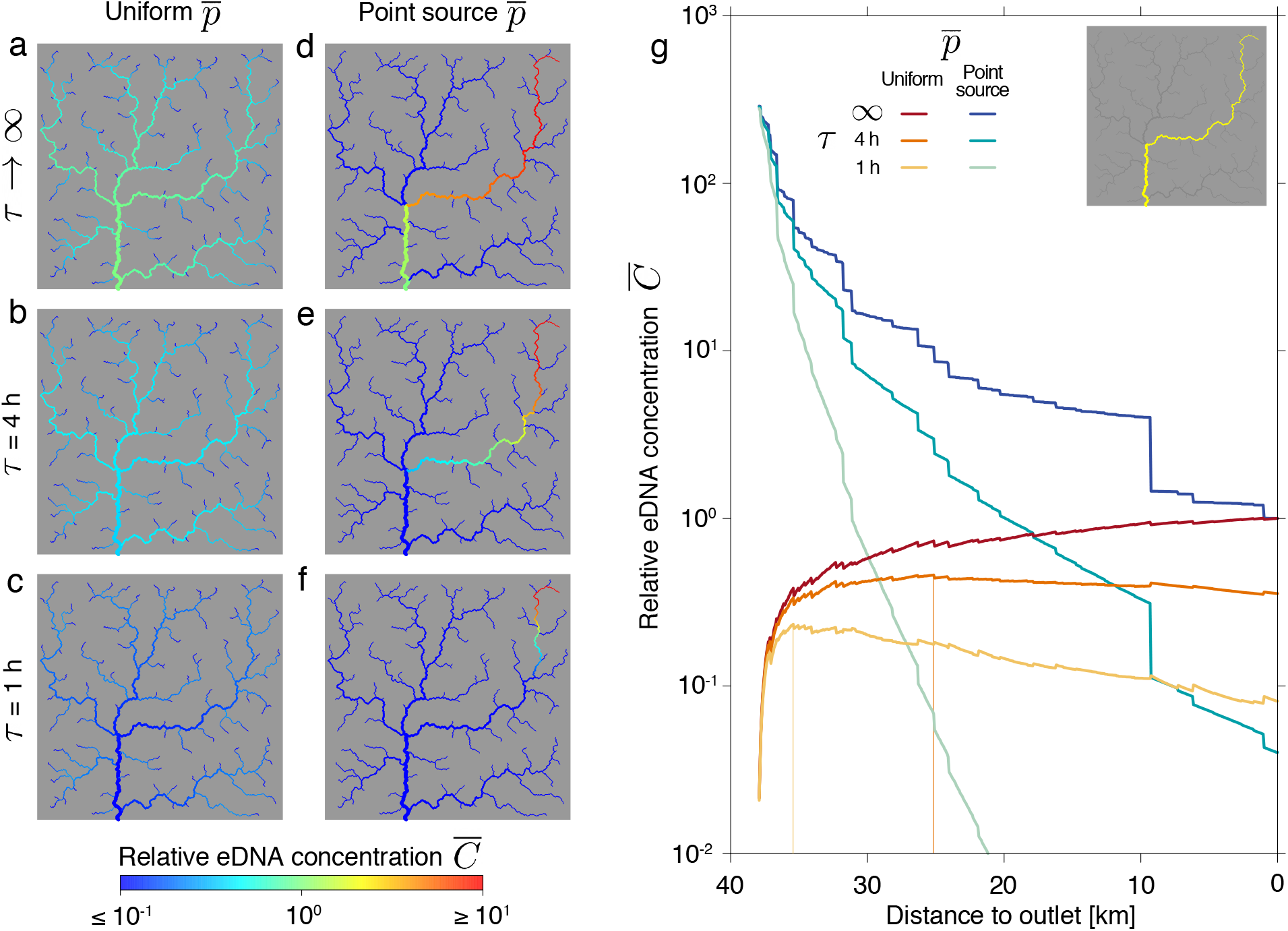
Effect of taxon distribution and eDNA decay processes on spatial patterns of eDNA concentration. a–f) Maps of relative eDNA concentration 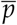 for uniform (panels a–c) and point-source (panels d–f) distributions of normalized taxon density 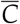 and no decay dynamics (panels a, d), *τ* = 4 h (panels b, e), *τ* = 1 h (panels c, f). In the point-source scenario, the taxon is concentrated in the pixel representing the headwater of the main river stem (i.e. the farthest point from the outlet). g) Profiles of 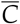 along the main river stem (depicted in yellow in the inset) for the six maps displayed in panels a–f. Vertical colored lines identify positions of maxima of 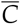 for profiles corresponding to maps on panels b and c.

The above-described pattern of increasing concentration in the downstream direction can be altered in the presence of eDNA decay. When 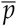 is uniform, the decay process induces a decreasing pattern of 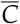 in the downstream direction in the region that is closer to the outlet, while, in the upstream reaches, 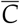 increases downstream for the aforementioned reasons (Figure 4b, c). As a result, the profile of concentration along the main stem is unimodal, and the location where 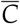 is maximized is controlled by the value of decay time (being shifted downstream for increasing values of *τ*, see vertical colored lines in Figure 4g).

When 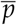 is concentrated in a single source and *τ* → ∞ (Figure 4d), eDNA concentration decreases downstream as a result of dilution because of water coming from “clean” tributaries. The concentration profile along the main stem (Figure 4g) is characterized by a number of vertical steps, which correspond to the confluences with the main tributaries. The inclusion of eDNA decay dynamics (Figure 4e, f) enhances the decreasing trend of 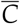, such that concentration values at the outlet are considerably lower than the corresponding values obtained with equal values of *τ* but uniform distribution for 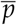.

### 3.2 Effect of sampling strategy

Aggregated results on the prediction skill expressed in terms of the presence/absence (*PA*) criterion (Eq. (7)) are shown in Figure 5. As expected, higher sampling intensity leads to better prediction skill, all other factors being equal. When the taxon is concentrated in few hotspots across the river network (S distributions) and only 10% of the sites are sampled (Figure 5a), the most reliable strategy is to preferentially sample the downstream sites (positioning U0), and the performance of the various strategies decreases with increasing percentage of sites sampled in the upstream region. Notably, with the aforementioned settings (i.e. S distribution, few sampling sites), this result holds regardless of the model variant. If the fraction of sampling sites increases (Figure 5b, c), the trend of increasing performance with increasing percentage of sampling sites in the downstream region holds only if perfect knowledge of the decay time and no measurement error are assumed (model variant TF); for the other model variants, more balanced strategies (U20 or U50) are to be preferred (Figure 5b), or even strategies that preferentially include upstream sites (U80) when the fraction of sampled sites is increased to 50% (Figure 5c, model ME).

**Figure 5:**
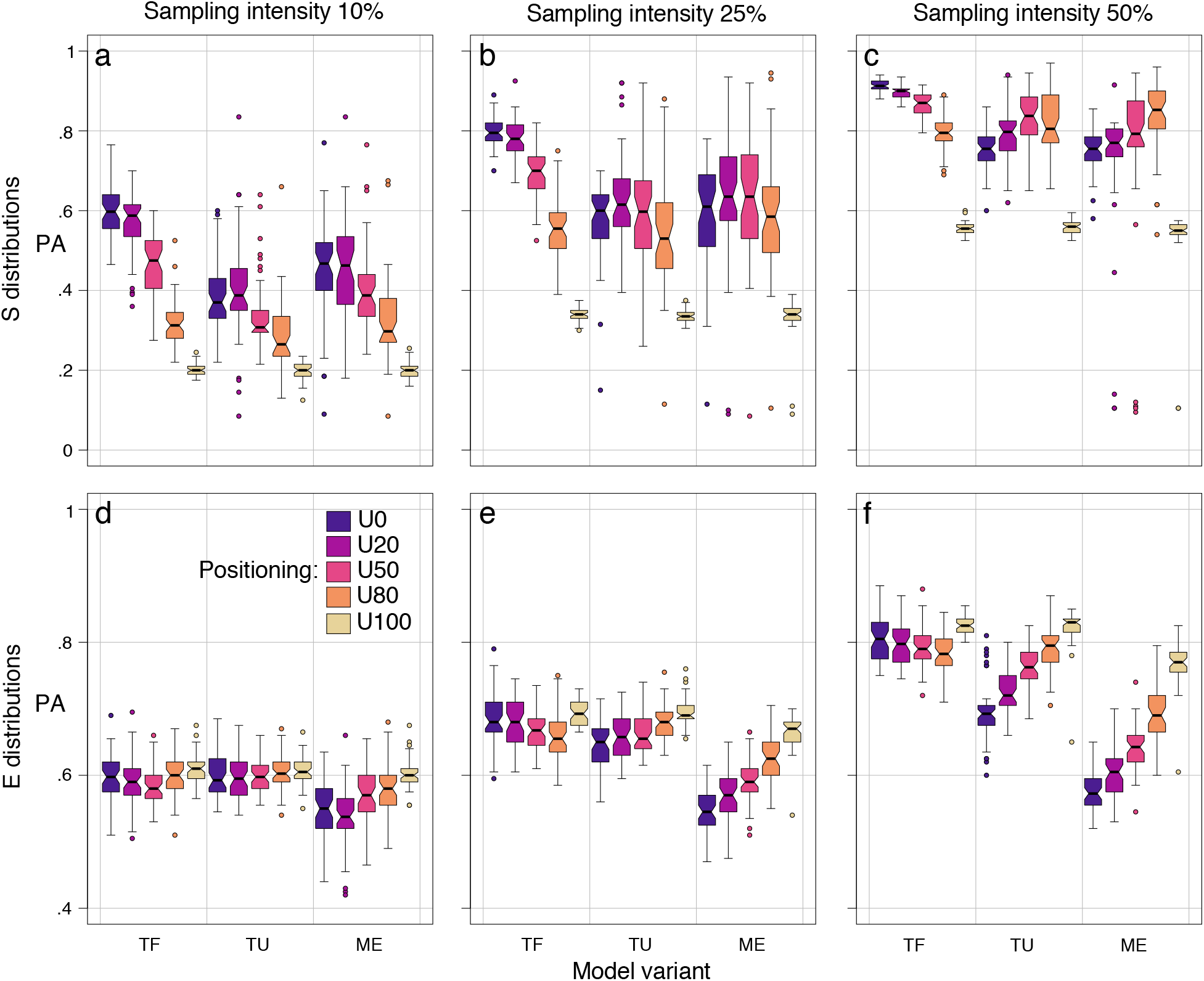
Boxplots of prediction skill expressed via the presence/absence (*PA*) criterion. Each boxplot is representative of 50 model runs (5 taxon distribution replicates times 10 sampling strategy replicates). Top and bottom rows refer to taxon distribution types S and E, respectively. Columns refer to different sampling intensities.

When instead the taxon is more evenly distributed across the catchment (E distributions - Figure 5d, e, f), the prediction skill is generally less sensitive to the sampling intensity and strategy: indeed, *PA* is above 0.4 with as few as 20 sampling sites (Figure 5d), for any strategy and model variant. Increasing sampling intensity to 50% only leads to mild improvements in prediction skill: *PA* is seldom (30.3% of the simulations) greater than 0.8 (Figure 5f), while this was more often the case (47.7% of the simulations) for the S distributions (Figure 5c). In general, for taxon distributions of the E type, the most convenient sampling strategy is to preferentially sample in the upstream region (U100), especially when the decay time is unknown and observed eDNA concentrations are affected by measurement error (model ME). The reason for such finding is that patterns of 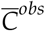 of type E are rather noisy, which makes it difficult for the eDITH model to disentangle the various contributions to the eDNA signal; preferentially sampling the upstream sites hence leads to improved predictions at the upstream reaches, since these signals are representative of few (or single) sources and thus easier to interpret.

Figure S2 shows prediction skill of the eDITH model expressed in terms of the density (*D*) criterion (Eq. (8)). Results are generally very similar to those discussed for the *PA* criterion. The main difference is that, while values of *D* are only slightly lower than the corresponding values of *PA* for taxon distributions of the S type (compare the top row of Figure S2 with the top row of Figure 5), the same is not true for taxon distributions of the E type: in this case, the fraction of sites where taxon density is correctly predicted is almost never (0.004% of the simulations) above 60% (Figure S2d, e, f), while the fraction of sites where the presence/absence status is correctly predicted is often (71.9% of the simulations) above 60% (Figure 5).

Remarkably, model variant TF generally performs better than other variants, all other factors and prediction skill criteria being equal. Instead, results obtained for model variant ME are not consistently worse than those obtained for variant TU (all other factors and prediction skill criteria being equal), despite the fact that, in the former model variant, observed concentrations 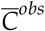 are affected by false negatives and measurement errors. In particular, when the taxon distribution is characterized by few hotspots, model variant ME performs similarly to (and, at times, better than) model TU (top rows of Figures 5 and S2). Conversely, considering false negatives and measurement errors worsens the prediction skill of the eDITH model when the taxon is more evenly distributed (bottom rows of Figures 5 and S2). To explain such finding, it has to be noted that, in the taxon distributions of type E, eDNA concentrations 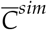 are often rather low (say, between 0.1 and 1, see Figure 3) and therefore possibly interpreted as false negatives, according to Eq. (6). Instead, in the S distributions, patterns of 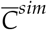 are characterized by few locations where the concentration is very high and unlikely to lead to a non-detection, resulting in observed values 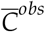 that are less perturbed. As a result, when *τ* is not known but the observed eDNA concentration 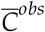 is rather high in some sites, the presence of a measurement error on 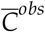 does not impact the estimates on taxon distribution performed by the eDITH model: indeed, the resulting 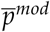 pattern will anyways resemble the real taxon distribution (at least to the same degree as if eDNA concentrations could be measured without error), with the possible drawback that the estimate of *τ* might be less accurate. As shown in Figure 6 in fact, values of the decay time estimated by the model variant ME are farther from the correct value *τ* = 4 h with respect to the TU model, and tend to coincide with the lower bound *τ* = 1 h imposed in the model fitting process, indicating failure in the determination of the value for such parameter.

**Figure 6:**
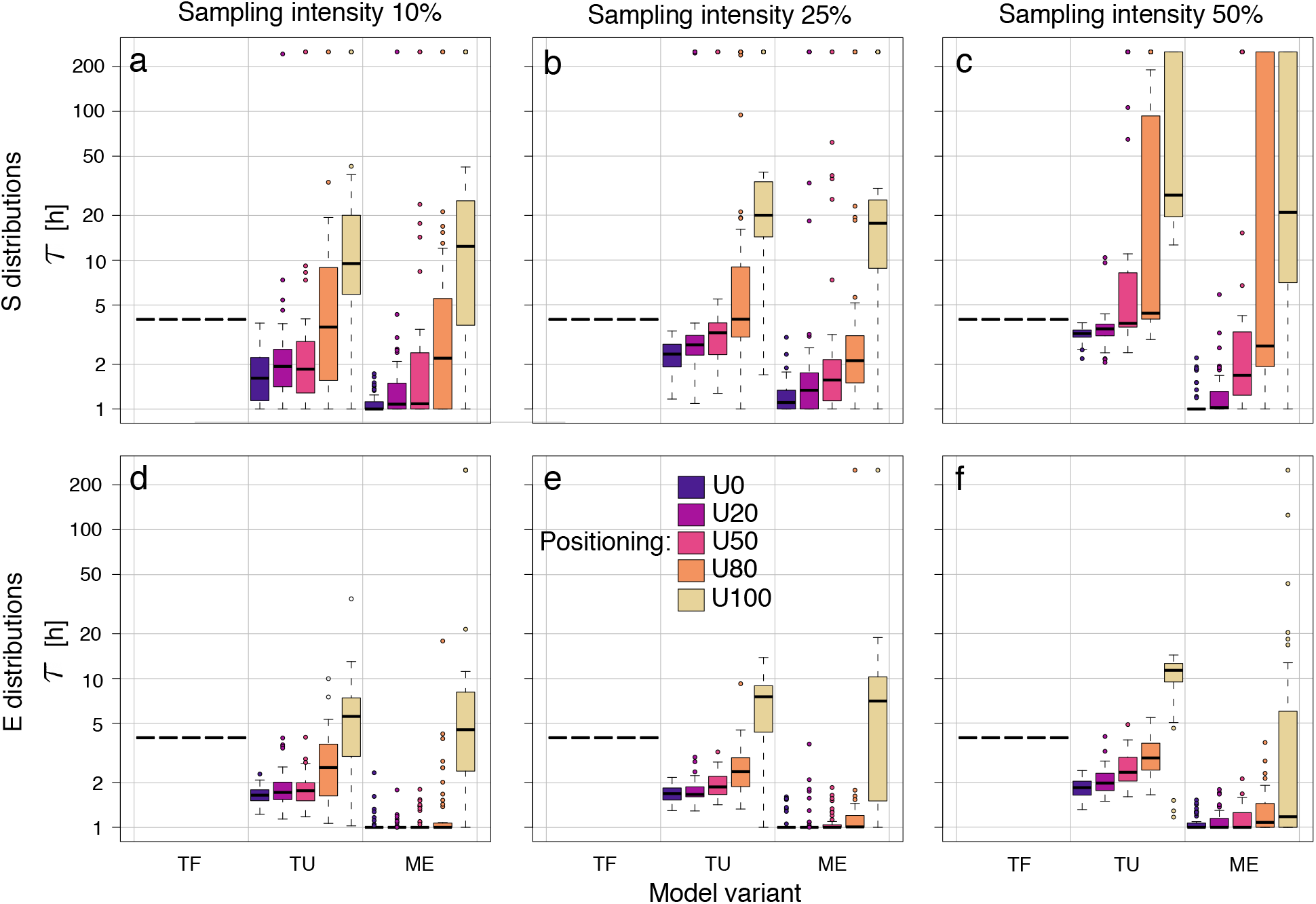
Boxplots of decay time values (in logarithmic scale) estimated by the model fitting procedure (step 4 of Figure 2). Symbols as in Figure 5.

As expected, an increase in sampling intensity leads to estimates of the decay time *τ* that are closer to the correct value *τ* = 4 h, all other factors being equal (Figure 6). In general, sampling strategies with preferential sampling at the downstream sites tend to lead to an underestimation of the decay time, while the opposite is true for sampling strategies with higher fraction of sites chosen among the upstream reaches (Figure 6). As a result, more balanced (e.g. U50) strategies might be more likely to accurately estimate the decay time dynamics. The fact that upstream-biased strategies tend to overestimate decay times (and, consequently, underestimate decay velocity) is explained by considering that these sampling schemes are characterized by a reduced degree of nestedness, namely the fraction of sampling sites that are connected by flow is low (details are provided in the Supporting Information). Indeed, in order to correctly assess decay dynamics, it is crucial to sample eDNA multiple times along the flow direction; if this were not the case, measurements from a single sample site that is not accompanied by other sampling sites upstream would make it impossible to disentangle the relative contributions of shedding (proportional to taxon density) and decay dynamics in the eDNA signal collected.

## 4 Discussion

Inferences of biodiversity based on riverine DNA data that go beyond the mere presence/absence status of organisms at the catchment level (i.e., gamma diversity only) need to consider hydro-morphological processes that control advection and decay of eDNA in stream water. In this work, we have exploited a hydrology-based eDNA transport model applied to a virtual—yet realistic—river network in order to analyze patterns of eDNA concentration across a river network and assess optimal strategies for eDNA sampling. Indeed, we found that spatial patterns of eDNA concentration are shaped by the scaling character of dendritic river networks: in particular, because of the power-law scaling of hydrological variables (chiefly river width), even a uniform taxon distribution does not lead to a uniform pattern of eDNA concentration, even if decay dynamics are neglected.

Our most important result is that selection of an optimal sampling strategy requires basic knowledge on the expected taxon distribution. When a taxon is expected to be concentrated in a few hotspots (e.g. as in the case of endangered and often only locally occurring organisms, such as the freshwater pearl mussel (*Margaritifera margaritifera*)) and sampling can be performed only at few sites due to time or budget constraints, it is advisable to preferentially sample the downstream locations; if, at a later stage, further sampling sites can be added, these should better be located in the upstream portion of the catchment. In such a case, the prediction skill of the eDITH model is quite sensitive to the number of sampling sites, and the fraction of sites where taxon density is correctly predicted does not change substantially with respect to the value related to correct presence/absence prediction. Conversely, when the taxon is distributed in a more uniform way across the catchment (e.g. as known for many widely distributed and common species, such as *Gammarus fossarum* or *Salmo trutta*), both number and positioning of sampling sites have a limited effect in improving the prediction skill of the model, and thus also the overall detection and resolution of biodiversity patterns of organisms across the catchment. Indeed, in such a case, values of observed eDNA concentration will tend not to differ much across sites, which complicates the localization of the main sources where eDNA is shed. However, we found that a preferential sampling of upstream sites might give slightly better overall predictions. Actually, in this case the improvement of prediction skill given by this strategy is limited to the upstream parts of the catchment, because the eDNA measurements collected therein would be representative of a lower number of sources and hence more easily interpretable by the eDITH model. In general, when the taxon is rather uniformly distributed, the presence/absence status can be fairly well predicted even with a limited number of sampling sites, while estimates of taxon density are much poorer than in the case when the taxon is located in few hotspots.

As expected, we observed that knowing the value of the decay time *τ* generally leads to improved prediction skill. Recently, research interest in the assessment of decay rates of eDNA collected from freshwater has expanded considerably [Eichmiller et al., 2016; Lance et al., 2017; Sassoubre et al., 2016; Seymour et al., 2018; Tsuji et al., 2017]. Such insights are of great help in improving the reliability of predictions of taxon distribution performed by the eDITH model, at least by allowing the determination of a narrow feasible range for this parameter. If instead *τ* is unknown, estimates of the taxon distribution operated by eDITH can still be fairly reliable, although the estimate of *τ* given by the model can be inaccurate. Measurement errors, which include all possible sources of uncertainties arising from the eDNA collection and sequencing processes, can indeed perturb the observed values of eDNA concentration, but they do not hinder the possibility to correctly estimate the taxon distribution, especially if the taxon is concentrated in a limited number of hotspots.

In order to perform a distinction between the various sampling strategies, we operated a partitioning of the reaches constituting the river network into upstream and downstream sites based on the distribution of drainage area values across the sites. It has to be noted that, owing to the criterion adopted, those marked as “downstream” sites do not only correspond to locations along the main stem and close to the river outlet, but also to sites that are rather distant from it (see Figure 1). Moreover, the so-obtained OCN has a drainage density of 0.7 km^−1^, which is representative of a very arid catchment (typical values of drainage density range from 2 to 20 km^−1^, with higher values corresponding to intermediate values of annual precipitation [Moglen et al., 1998]). We refrained from using an OCN with higher drainage density because this would have implied increasing the dimensionality of the problem, which would have hindered the convergence of the calibration algorithm, and likely jeopardized the prediction skill of the model. This implies that the headwater streams shown in Figure 1 (marked with red dots, and corresponding to the “upstream” sites of the computer simulations) are not likely to correspond to reaches of Strahler stream order equal to 1 of a real catchment of the same size, but rather to larger reaches, with Strahler order values of at least 2. Consequently, sites here marked as “downstream” are (roughly) representative of real stream reaches with Strahler order greater than 2.

Importantly, to perform the computer simulations, we partitioned the river network at the reach scale, thereby implicitly assuming that sampling more than once along a single river reach is sub-optimal, because this would lead to measurements whose information content is overlapping. In general, we suggest to choose sampling sites such that their pairwise along-stream distance is of at least a couple of kilometers (for instance, in this application we fixed the maximum reach length equal to 2.5 km), so that they are representative of subcatchments that are rather distinguished. However, in the presence of high stochasticity (and/or high likelihood of non-detection) of measured eDNA concentrations within a single site, it might be relevant to perform multiple sampling at the same site. Potentially, the number of samples taken at a site might change with the site’s position across the river network (e.g. more samples per site taken at the downstream locations, as suggested by Bylemans et al. [2018]). A possible development of our modelling work could actually verify this hypothesis. Collecting more samples at fixed downstream locations would also be beneficial because, due to a dilution effect, eDNA concentrations at the downstream sites are likely not high even in the presence of a local taxon hotspot (see Figure 3), and hence more prone to result in non-detections. This is especially important in the context of many metabarcoding studies setting thresholds of read numbers below which the signal is no longer accepted as a true positive [Deiner et al., 2017; Mächler et al., 2020]. In this perspective, collecting larger water volumes at the downstream sites is likely to help minimize the risk of non-detection.

In this study, for the sake of simplicity, we focused on localization of a single taxon within a river basin. However, the eDITH model herein adopted can also be applied repeatedly with eDNA data for different taxa collected at the same sites (also in the form of metabarcoding data, i.e. read numbers) in order to estimate spatial patterns of biodiversity [Carraro et al., under review]. Hence, the findings of the present study can directly be generalized to the issue of determining the optimal sampling strategy within a river network in order to maximize information on the spatial distribution of biodiversity.

In summary, our study revealed potential strategies to optimize the design of eDNA sampling campaigns in river networks. Notably, the insights gained by our modelling approach need be paired by biological and empirical knowledge on the investigated taxon: in fact, the choice of the optimal sampling design relies on preliminary knowledge on the expected distribution of the taxon, and on whether or not estimates of eDNA decay rates in the environment are available. Coupling modelling and empirical evidence on eDNA transport and decay processes is thereby crucial in order to fully exploit the power of eDNA to monitor freshwater biodiversity.

## Author contributions

LC and FA developed the idea, LC conceptualized the work and wrote the first draft of the manuscript. LC and JBS analyzed the data. All authors contributed to the final manuscript version.

## Acknowledgments

FA acknowledges support from the Swiss National Science Foundation grants No. PP00P3 179089 and 31003A 173074 as well as from the University of Zurich Research Priority Programme “URPP Global Change and Biodiversity”. We thank Jeanine Brantschen and Rosetta C. Blackman for valuable comments on the manuscript.

## Notes

### Competing Interest Statement

The authors have declared no competing interest.

