## Supporting Information for "How to design optimal eDNA sampling strategies for biomonitoring in river networks"

#### Supporting Text

##### Variation of $\overline{C}$ across a river network when $\overline{p}$ is uniform

From Eqs. (1) and (4), when $\overline{p}_i = \overline{p}\ \forall i = 1, \ldots, N$ and $\tau \to \infty$, one gathers

$$\frac{\overline{C}_j}{\overline{p}} = \frac{\left(\sum_{i\in\gamma(j)} A_{S,i}\right)/Q_j}{\left(\sum_{i\in\gamma(o)} A_{S,i}\right)/Q_o}. \tag{S1}$$

Note that the denominator of Eq. (S1) does not depend on $j$. Moreover, it is $Q_j \sim A_j$, with $A_j = N_j L_P^2$, where $N_j$ is the number of pixels upstream of $j$ (including $j$ itself) and $L_P$ the length of a pixel (corresponding to 50 m in this application); $A_{S,i} = L_i w_i$, with $L_i = L_P$ or $L_i = L_P\sqrt{2}$ if flow direction is horizontal/vertical or diagonal, respectively; $w_i \sim A_i^{0.5}$ (see Eq. (2)). If, for the sake of simplicity, we assume that all flow directions are horizontal or vertical (generalization in the case of a fraction of the flow directions being diagonal is straightforward), Eq. (S1) becomes

$$\frac{\overline{C}_j}{\overline{p}} \sim \frac{\sum_{i\in\gamma(j)} A_i^{0.5}}{A_j} \sim \frac{\sum_{i\in\gamma(j)} N_i^{0.5}}{\sum_{i\in\gamma(j)} N_i^{0}} \tag{S2}$$

where the r.h.s. of Eq. (S2) is obtained by considering that $A_j \sim N_j = \sum_{i\in\gamma(j)} N_i^0$. From Eq. (S2), the ratio $\overline{C}_j/\overline{p}$ would be independent of $j$ only if the numerator of the r.h.s. were equal to $\sum_{i\in\gamma(j)} N_i^0$ (i.e., if the exponent for the $w \sim A$ scaling of Eq. (2) were equal to 0, which would imply constant width in the downstream direction). Actually, the numerator of the r.h.s. grows faster than the denominator in the downstream direction (because $N_i \geq 1\ \forall i = 1, \ldots, N$, and $N_j > N_i$ if $j$ is downstream of $i$), and therefore the ratio $\overline{C}_j/\overline{p}$ increases in the downstream direction.

##### Nestedness of a sampling design

The degree of nestedness of a set of sampling sites can be evaluated as the number of pairs of sites that are connected by flow. In order to allow comparison between sampling designs with a different number of sites $N_S$, the value calculated above can be normalized by dividing it by its maximum value $\sum_{i=1}^{N_S-1} i$, corresponding to a design where all sites are connected by a single flow path. The so-obtained values of normalized nestedness are displayed in Figure S3. Sampling designs with larger fraction of sites from the downstream region of the catchment are characterized by larger values of nestedness, with differences in positioning becoming more significant with increasing sampling intensity.

### Supporting Tables

**Table S1:** List of symbols used in the manuscript.

| Symbol | Description | Dimension |
|---|---|---|
| $A_i$ | Drainage area at node $i$ | $L^2$ |
| $A_{S,i}$ | Source area at node $i$ | $L^2$ |
| $A_T$ | Threshold value of drainage area used to distinguish the perennial river network | $L^2$ |
| $C_i$ | eDNA concentration at node $i$ | $NL^{-3}$ |
| $\overline{C}_i$ | Normalized eDNA concentration at node $i$ | - |
| $\overline{C}_i^{obs}$ | Observed (step 3 of Figure 2) normalized eDNA concentration at node $i$ | - |
| $\overline{C}_i^{sim}$ | Simulated (step 2 of Figure 2) normalized eDNA concentration at node $i$ | - |
| $D$ | Prediction skill index based on density | - |
| $K_i$ | Natural logarithm of $\overline{C}_i^{obs}$ | - |
| $L_i$ | Length of the reach corresponding to node $i$ | L |
| $L_{ij}$ | Length of the path connecting node $i$ to $j$ | L |
| $L_P$ | Length of a pixel side | L |
| $N$ | Number of nodes into which the river network is partitioned | - |
| $N_i$ | Drainage area at node $i$ (in number of pixels) | - |
| $N_H$ | Number of hotspots of taxon density | - |
| $N_S$ | Number of sampling sites | - |
| $p_i$ | eDNA production rate | $NL^{-2}T^{-1}$ |
| $\overline{p}_i$ | Normalized eDNA production rate | - |
| $\overline{p}_i^{mod}$ | Modelled (step 4 of Figure 2) normalized eDNA production at node $i$ | - |
| $\overline{p}_i^{sim}$ | Simulated (step 1 of Figure 2) normalized eDNA production rate at node $i$ | - |
| $\overline{p}_i'$ | Temporary (non-normalized) eDNA production rate | - |
| $Q_i$ | Water discharge at node $i$ | $L^3T^{-1}$ |
| $v_i$ | Water velocity at node $i$ | $LT^{-1}$ |
| $v_{ij}$ | Average (in space) water along the path connecting node $i$ to $j$ | $LT^{-1}$ |
| $w_i$ | River width at node $i$ | L |
| $\gamma(i)$ | Set of nodes upstream of $i$ (including $i$) | - |
| $\tau$ | eDNA decay time | T |

### Supporting Figures

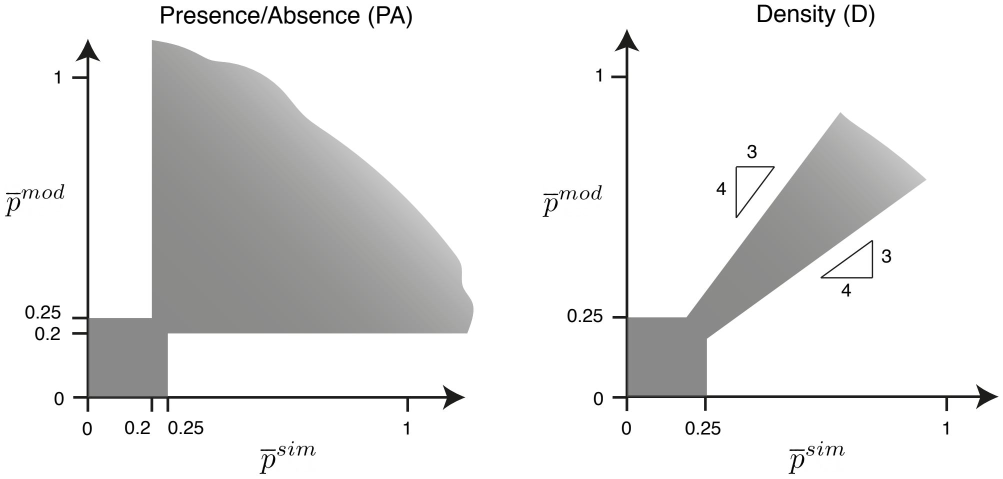


**Figure S1:** Graphical representation of the two prediction skill criteria used. If, for a given site, the pair $\left[\overline{p}^{mod}, \overline{p}^{sim}\right]$ falls within the gray region, then that site is deemed correctly predicted.

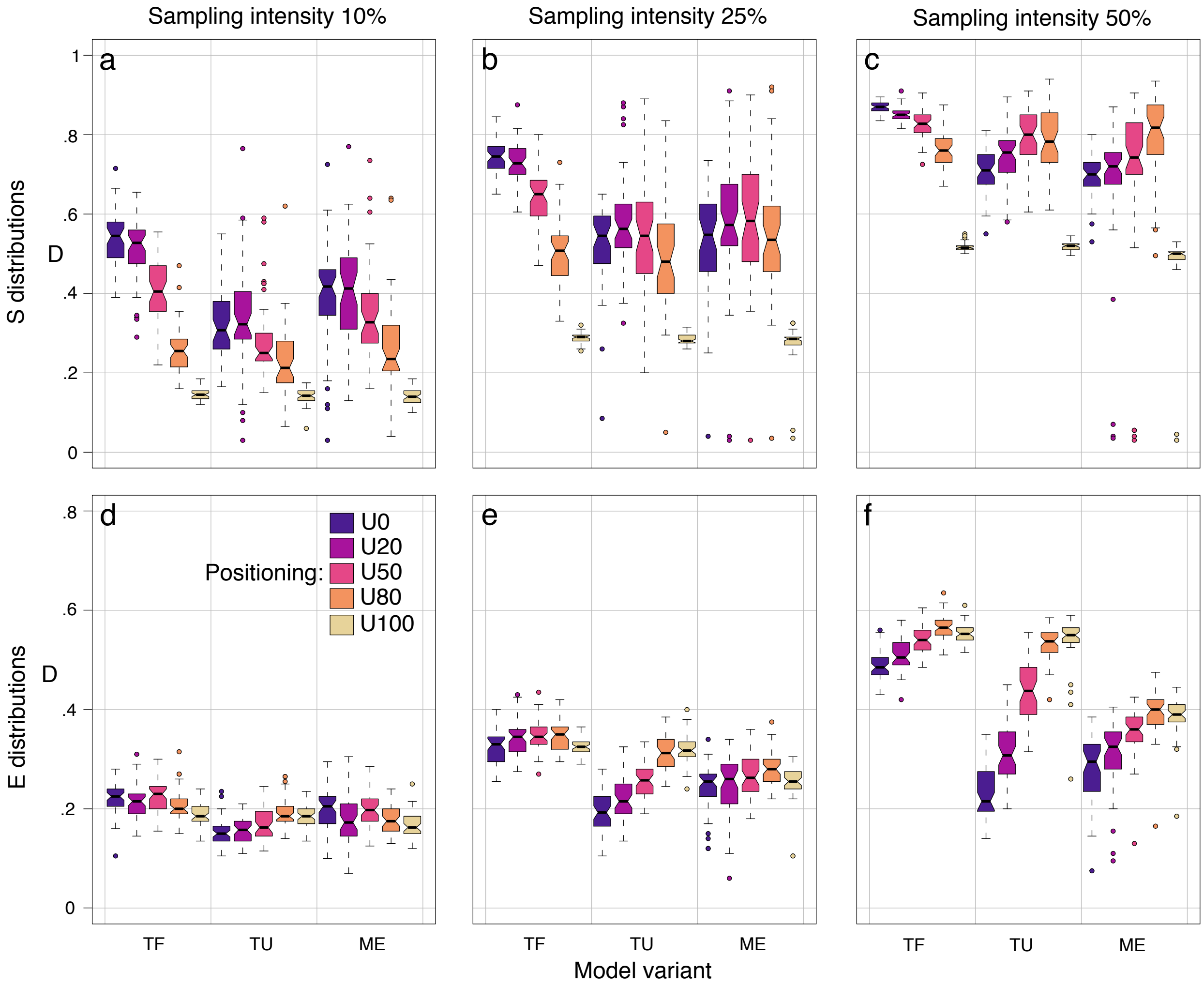


**Figure S2:** Boxplots of prediction skill expressed via the density (*D*) criterion (see Eq. (8)). Symbols as in Figure 5.

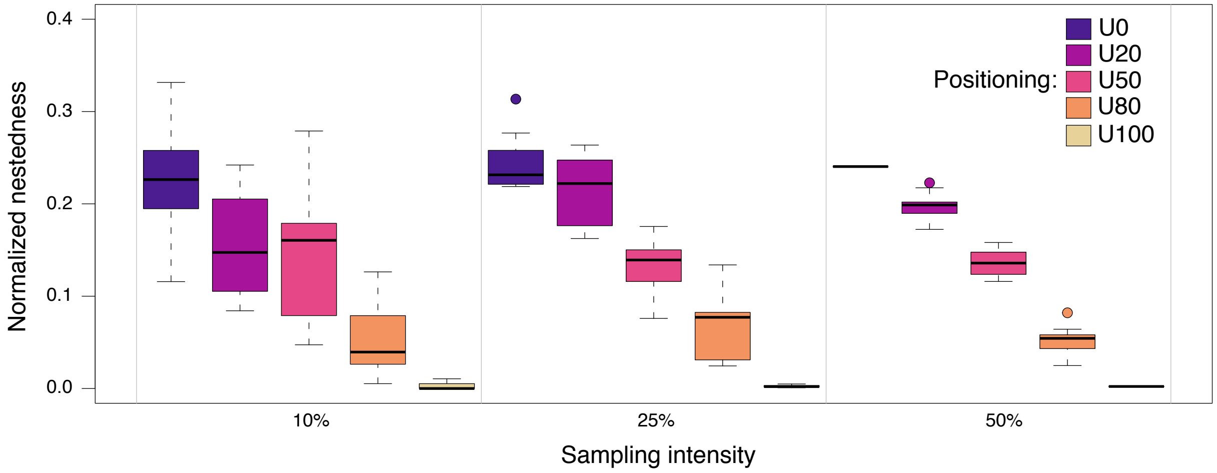


**Figure S3:** Boxplots of normalized nestedness for the different sampling designs. Each boxplot is representative of 10 values.
